# Age-dependent decrease of hepatic geranylgeranoic acid and its oral supplementation prevents spontaneous hepatoma in C3H/HeN mice

**DOI:** 10.1101/2021.04.16.440102

**Authors:** Yuki Tabata, Masahide Omori, Yoshihiro Shidoji

## Abstract

**Aim:** Geranylgeranoic acid (GGA) has been developed as a preventive agent against second primary hepatoma. Recently, GGA was reported to induce cell death in human hepatoma cells via TLR4-mediated pyroptosis. We have reported that GGA is enzymatically biosynthesized from mevalonic acid in human hepatoma-derived cells and that endogenous GGA is found in most organs of rats. In addition, we found that upregulation of endogenous GGA levels by zaragozic acid A (ZAA) induced cell death in human hepatoma-derived cells. Therefore, we investigated the age-related changes of hepatic GGA and the possibility of suppressing hepatocarcinogenesis by GGA supplementation using male C3H/HeN mice that spontaneously develop hepatoma.

**Methods:** We measured endogenous GGA and mRNA of MAOB, a key enzyme of GGA biosynthesis, in the liver of male C3H/HeN mice aged 6–93 weeks. We also tried suppressing spontaneous hepatocarcinogenesis by a single administration of GGA to C3H/HeN mice.

**Results:** Hepatic GGA content and *Maob* mRNA expression level age-dependently decreased in male C3H/HeN mice, some of which produced spontaneous hepatoma in 2 years. A single oral administration of GGA at 11 months of age significantly prevented hepatoma in terms of the number and weight of tumors per mouse at 23 months. Oral supplementation with GGA or geranylgeraniol significantly increased endogenous hepatic GGA contents dose-dependently, and ZAA dramatically upregulated hepatic GGA.

**Conclusion:** In this study, we found an age-dependent decrease in hepatic endogenous GGA in male C3H/HeN mice and efficient prevention of spontaneous hepatoma by a single administration of GGA at 11 months of age.

## INTRODUCTION

We have been studying the preventive effect of retinoids against hepatoma. In the course of these studies, Muto and Moriwaki *et al*. proposed the concept of “acyclic retinoids” based on their study of the relationship between the ligand activity of retinoid receptors and the induction of hepatoma differentiation^[1,2]^. In other words, acyclic retinoids have ligand activity against intracellular retinoic acid-binding proteins (CRABP1 and CRABP2) and retinoid receptors (RAR and RXR), and induce differentiation of hepatoma cell lines to enhance albumin production. In fact, when we checked whether acyclic retinoids have the same “retinoid action” as all-*trans*-retinoic acid (ATRA, whose chemical structure is shown in **Figure 1A**), we found that, one acyclic retinoid significantly induced the expression of neurotrophic receptor kinase 2 (NTRK2) and RARβ, as well as ATRA, in the human neuroblastoma cell line SH-SY5Y ^[3]^.

**Figure 1.**
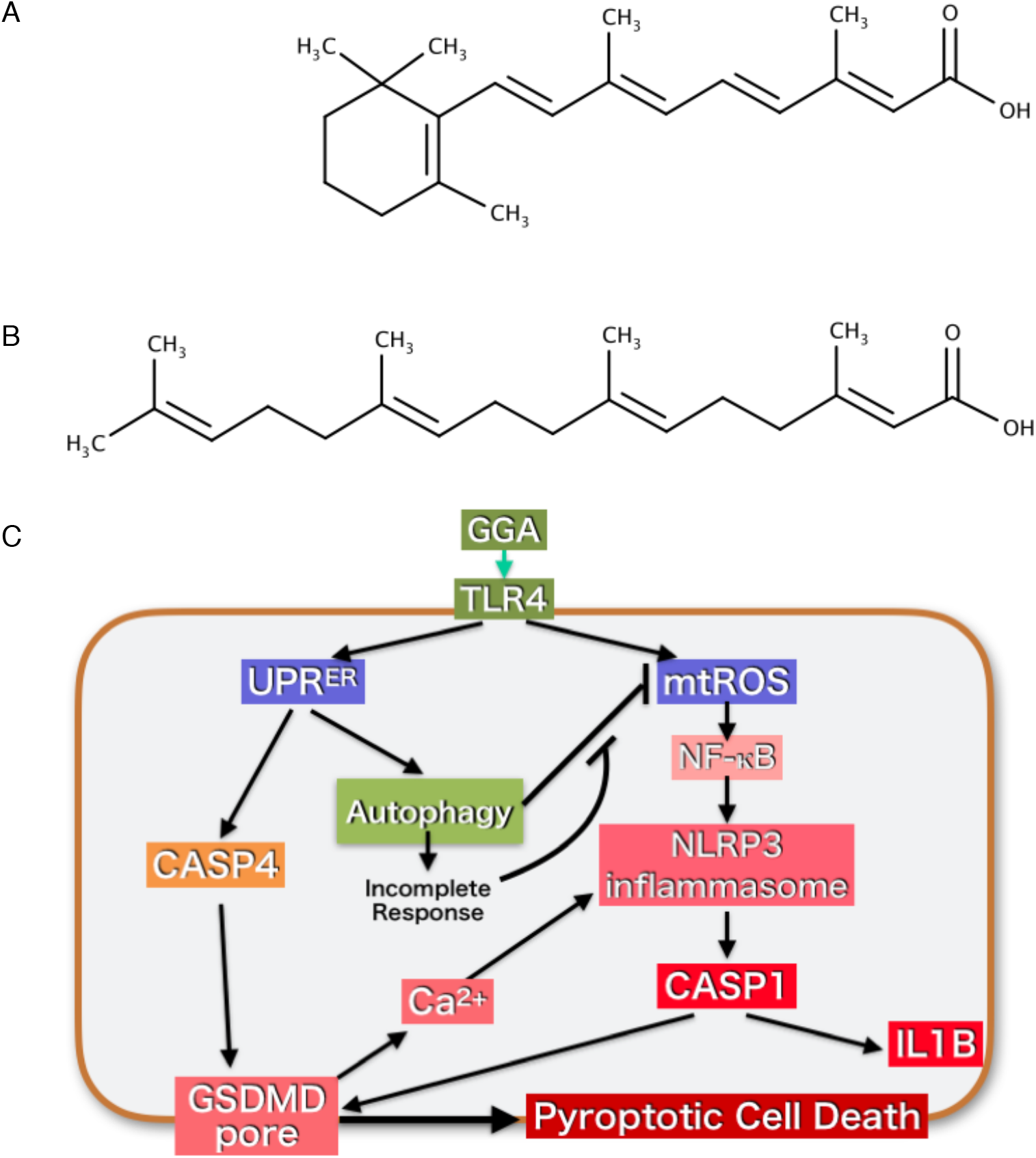
The chemical structure of geranylgeranoic acid (GGA) and proposed mechanism of GGA-induced cell death in human hepatoma cells. A: The chemical structure of all-*trans* retinoic acid (ATRA), which is a natural monocyclic retinoid. B: The GGA contains three *trans*-*cis* isomeric double bonds, and although there are reports of 2-*cis* isomers as metabolites among the stereoisomers, here we show only the structure of the 2,6,10-*trans* isomer. C: In human hepatoma-derived cell lines, GGA has been reported to induce an endoplasmic reticulum stress response (UPR^ER^)^[11]^, an incomplete response of autophagy^[10]^, activation of CASP4 and cleavage of gasdermin D (GSDMD), activation of CASP1 and further cleavage of GSDMD and activation of the NLRP3 inflammasome^[12]^. In this context, TLR4 signaling-mediated pyroptosis has been proposed as a mechanism of GGA-induced cell death^[12]^.

Among acyclic retinoids so far investigated, geranylgeranoic acid (all-*trans* 3,7,11,15-tetramethyl-2,6,10,14-hexadecatetraenoic acid or GGA, **Figure 1B**) and its didehydro derivative (all-*trans* 3,7,11,15-tetramethyl-2,4,6,10,14-hexadecapentaenoic acid or peretinoin) have been developed as preventive agents against second primary hepatoma^[4,5]^. Indeed, the 4,5-didehydro derivative of GGA potentially prevented second primary hepatoma in placebo-controlled randomized phase II–III clinical trials^[6,7]^. Using some human hepatoma-derived cell lines, we found that both GGA and 4,5-didehydroGGA induced tumor-specific cell death, which may be involved in the suppression of second primary hepatocarcinogenesis^[8,9]^. As for the molecular and cellular mechanisms of GGA-induced cell death, several studies have so far been conducted. In these studies, GGA was reported to induce cell death in human hepatoma cells via endoplasmic reticulum stress response and incomplete autophagic response^[10,11]^. Most recently, we have reported that GGA induces Toll-like receptor 4 (TLR4)-mediated pyroptosis in human hepatoma-derived cells^[12]^.

The cell death-inducing effect of GGA on hepatoma cells, as shown on the left side of **Figure 1C,** is signaled by the cell surface receptor TLR4, which causes activation of CASP4 via the endoplasmic reticulum stress response or unfolded protein response, UPR^ER^, resulting in cleavage of gasdermin D (GSDMD) and the translocation of its N-terminal fragment to the plasma membrane leading to the formation of the GSDMD pore. Another TLR4 signaling, as shown on the right side of **Figure 1C**, causes increased reactive oxygen species (ROS) production in the mitochondria, leading to the transfer of NF-κB to the nucleus and promoting the formation of the NLRP3 inflammasome. In addition, the influx of extracellular Ca^2+^ causes inflammasome activation, which in turn activates CASP1, leading to extensive disruption of the cell membrane and completing cell death by pyroptosis. According to a recent review by Takahama, Akira and Saitoh^[13]^, autophagy also targets and inactivates the inflammasome, so we believe that the incomplete response of autophagy by GGA may support the activation of CASP1 by preventing the inactivation of the inflammasome.

We have previously reported that GGA is a natural acyclic diterpenoid present in some medicinal herbs^[14]^. Not only in plants, but also in animal cells, GGA was recognized as a mevalonic acid (MVA)-derived metabolite first in cell-free homogenates of the bovine retina in 1983^[15]^, and then in a parasitic worm in 1993^[16]^. Recently, we reported that GGA is endogenously present in various organs of male Wistar rats^[17]^. In particular, the liver contained 5 to 10-fold times more GGA than other organs. Biosynthesis of GGA from MVA via geranylgeranyl diphosphate is also confirmed in human hepatoma-derived cells [**Figure 2**]. We also reported that monoamine oxidase B (MAOB) is primarily involved in the oxidation of geranylgeraniol (GGOH) to geranylgeranyl aldehyde, which is a direct precursor of GGA^[18]^. Of note, a strong correlation was detected between endogenous GGA and *MAOB* mRNA levels in human hepatoma cells^[18]^.

**Figure 2.**
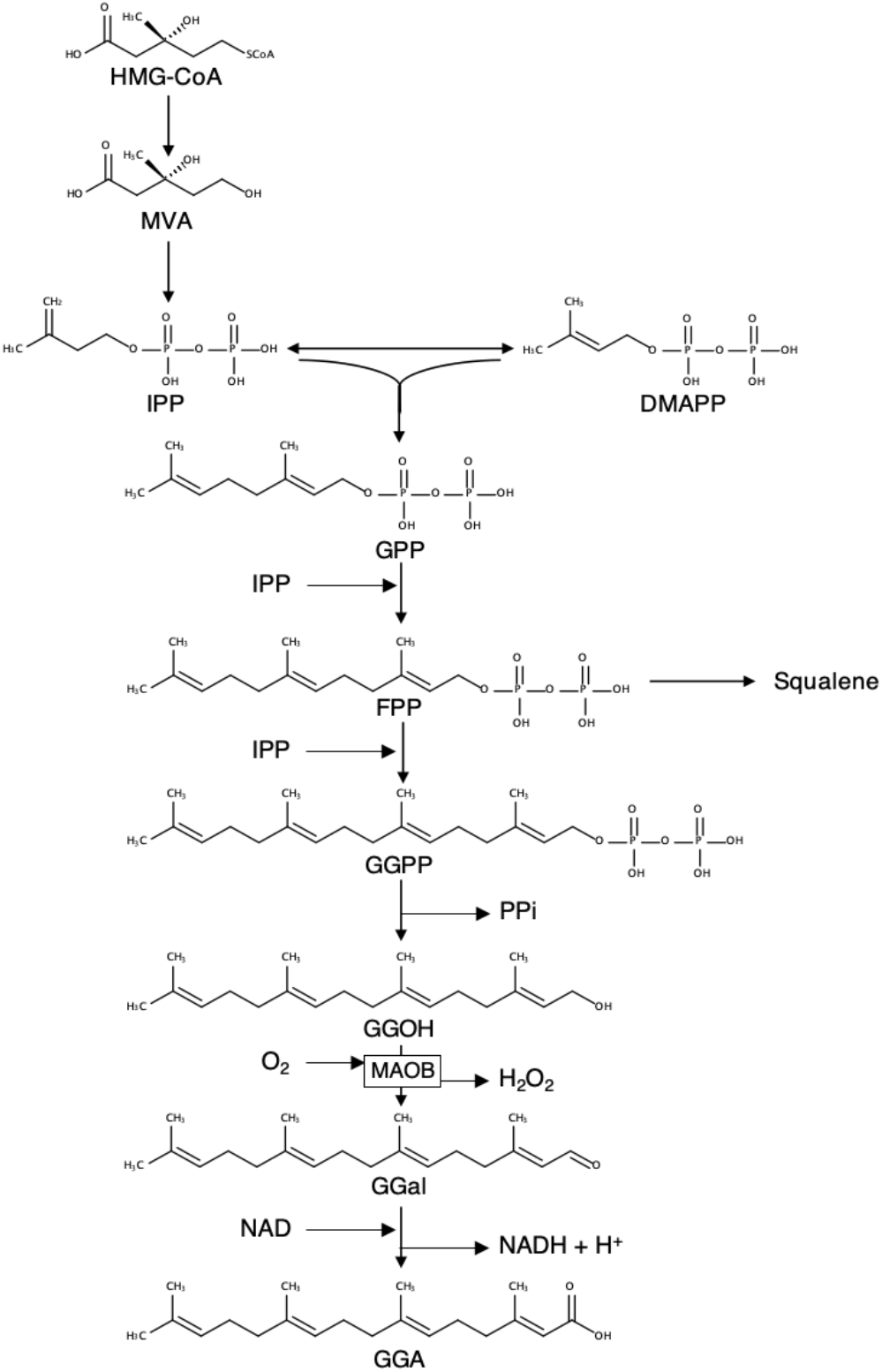
Metabolic pathway of GGA biosynthesis from HMG-CoA in human hepatoma cells. Hepatic biosynthesis of GGA from MVA was demonstrated in human hepatoma-derived HuH-7 cells by isotopomer spectral analysis^[17]^, and hepatic MAOB was shown to be involved in oxidation of GGOH to GGal, a key enzymatic step of GGA biosynthesis^[18]^. HMG-CoA: 3-hydroxy-3-methylglutaryl-coenzyme A; MVA: mevalonic acid; IPP: isoprenyl diphosphate; DMAPP: dimethylallyl diphosphate; GPP: geranyl diphosphate; FPP: farnesyl diphosphate; GGPP: geranylgeranyl diphosphate; GGOH: geranylgeraniol; GGal: geranylgeranyl aldehyde; GGA: geranylgeranoic acid; MAOB: monoamine oxidase B.

It has been well established that the human brain from postmortem samples shows an age-related increase in MAOB activity^[19]^. However, by using enzyme autoradiography with selective inhibitors, a previous study reported that the MAOB enzyme levels in mouse liver decreased with aging, whereas most other organs such as the brain, heart and kidney inversely increased the enzyme level along with aging^[20]^. Hence, we reasonably speculate that with age, *Maob* mRNA in mouse liver will decrease and correlated with this decrease, hepatic GGA level will also decrease with age.

In this context, we focused on male C3H/HeN mice, in which 40–100% of the mice spontaneously develop primary hepatoma after 2 years of normal rearing^[21,22]^. Therefore, C3H/HeN mouse is often used for experiments as a model animal of hepatic carcinogenesis^[23,24]^, although the mechanism underlying the hepatocarcinogenesis of this mouse has not yet been elucidated. Assuming that the putative age-dependent decrease in hepatic *Maob* mRNA level in C3H/HeN mice may lead to an age-dependent decrease in hepatic GGA level, which may allow an uncontrolled spontaneous development of primary hepatoma, supplementation of GGA at a time when hepatic GGA level of C3H/HeN mice is reduced may effectively suppress the spontaneous development of primary hepatomas. Indeed, in clinical trials^[4–7]^, taking 4,5-didehydroGGA for 1 year after a radical treatment of primary hepatoma significantly reduced the incidence of second primary hepatoma five years later.

In this study, we explored the relationship between hepatic GGA level in C3H/HeN mice and spontaneous hepatocarcinogenesis. First, we present the presence of endogenous GGA in the liver of C3H/HeN mice and age-dependent decrease of hepatic GGA level, which is associated with reduced expression of hepatic *Maob* mRNA. Next, we examined whether spontaneous hepatocarcinogenesis was avoided by oral administration of exogenous GGA. In addition, we recently reported that upregulation of endogenous GGA levels by zaragozic acid A (ZAA) induced cell death in human hepatoma-derived cells^[17]^. Hence, finally, we investigated dietary supplementation of GGA or GGOH, and intraperitoneal administration of ZAA to C3H/HeN mice to increase hepatic GGA content. The results obtained in this study suggest that the age-dependent decrease of endogenous GGA in the liver of C3H/HeN mice may be involved in spontaneous hepatocarcinogenesis, and the reason why oral administration of exogenous GGA avoided hepatocarcinogenesis.

## MATERIALS AND METHODS

### Chemicals

GGA was prepared by Kuraray Co. (Okayama, Japan) and Kowa Pharmaceutical (Tokyo, Japan). GGOH was provided by Eisai Foods (99% pure; Tokyo, Japan). Acetonitrile (LC/MS grade) and ethanol were purchased from Sigma-Aldrich (St. Louis, MO). Methanol was from Wako Pure Chemical Industries (Osaka, Japan). Chloroform was obtained from Kanto Chemical Co. (Tokyo, Japan). ZAA (an inhibitor of squalene synthase) was provided by Merck (Darmstadt, Germany). All chemicals other than those stated above were of reagent grade.

### Animals

Male C3H/HeN mice were purchased from Charles River Japan (Kanagawa, Japan) or CLEA Japan (Tokyo, Japan). The mice were fed a conventional diet (CE-2; CLEA Japan) and tap water. The animals (5 mice/group) were euthanized under isoflurane anesthesia by collecting blood through the vena cava and the liver was immediately excised, and instantaneously frozen at –25°C to determine endogenous GGA levels at each age from 6- to 93-week old. At that time, a part of the tissue was treated with RNA*later* (Thermo Fisher, Tokyo, Japan) and stored at room temperature until it was used for total RNA preparation, described below.

### Evaluation of inhibitory effects of GGA on the development of spontaneous hepatomas in C3H/HeN mice

After one week of normal rearing, the mice were divided into four groups (10 mice/group), GGA (50 μg/mouse) in peanut oil was orally administered to each group of mice through a gastric tube only once at 7, 11, and 17 months of age, respectively. The control mice were treated with the vehicle alone at 17 months of age. At 23 months of age, all mice were subjected to laparotomy under anesthesia to observe the presence of hepatomas with the naked eye, and their weight and number were measured after excision.

### Preparation of GGA- or GGOH-containing diets and feeding to C3H/HeN mice

Aliquots of GGA or GGOH (0–800 μg or 0–8 mg, respectively, dissolved in 300 μL ethanol) were dropped and soaked into each pellet (approximately 3 g) of CE-2 and the vehicle solvent was removed by overnight air ventilation in a draft chamber. After removal of the solvent, the pellets were kept frozen at –25 °C until use. These diets were fed to 24-week old male C3H/HeN mice at doses of GGA (0–8 μg/day/mouse) or GGOH (0–8 mg/day/mouse) for 1 week. After 1-week treatment with the experimental diets, the mice were sacrificed, and the liver was excised to measure hepatic endogenous GGA contents.

### ZAA administration to male C3H/HeN mice

Twenty-four-week old male C3H/HeN mice were treated with intraperitoneal injections of ZAA at a dose of 0.2 mg/kg B.W. After 0 to 12 h from the injection of ZAA, the mice were sacrificed, and the liver was excised to measure the endogenous hepatic GGA contents.

### RT-qPCR

Total RNA was prepared from mouse tissue using the Fastgene™ RNA Basic kit and DNase I (Nippon Genetics, Tokyo, Japan). For cDNA synthesis, Fastgene™ Scriptase II (Nippon Genetics) was used according to the manufacturer’s instructions. Real-time PCR was performed using LightCycler FastStart DNA Master^PLUS^ SYBR Green I (Roche Diagnostics, Tokyo, Japan) on a LightCycler 96 (Roche). Relative gene expression levels of *Maoa* and *Maob* were analyzed using the 2^−ΔΔ^Ct method with a reference of *28S rRNA*. Primer sequences and real-time PCR settings used in this study are presented in **Supplementary Tables 1** and **2**.

### Lipid extraction and quantitative measurement of GGA contents

Extraction of total lipids and LC/MS/MS quantification of GGA and arachidonic acid (ARA) in C3H/HeN mouse liver was performed using the procedures described in our previous study^[17]^.

### Statistical analysis

Statistical comparisons were performed using ANOVA with post hoc Scheffe test where appropriate. The SPSS statistical software package (IBM, Tokyo, Japan) was used for the analyses.

### Ethical approval

This research was approved by the Animal Experimental Ethics Committee of the University of Nagasaki (No. 29–29).

## RESULTS

### Endogenous hepatic GGA content of C3H/HeN mice decreases with age

First, we examined whether the amount of endogenous GGA in C3H/HeN mouse liver varied along with aging. Endogenous GGA levels in C3H/HeN mouse liver began to decrease after 20 weeks of age and remained significantly low from 25 to 60 weeks of age, and then declined rapidly until hepatic GGA became completely undetectable at 93 weeks of age [**Figure 3A**]. In other words, hepatic GGA level showed a two-step decrease with age. In the first step, hepatic GGA level in the male C3H/HeN mice decreased by approximately 30% from 18 to 25 weeks of age, and in the second stage, it decreased rapidly from 60 to 93 weeks of age and became undetectable. In contrast to GGA, hepatic content of free ARA, which is a structural isomer of GGA, rather increased with age [see **Supplementary Figure 1**].

**Figure 3.**
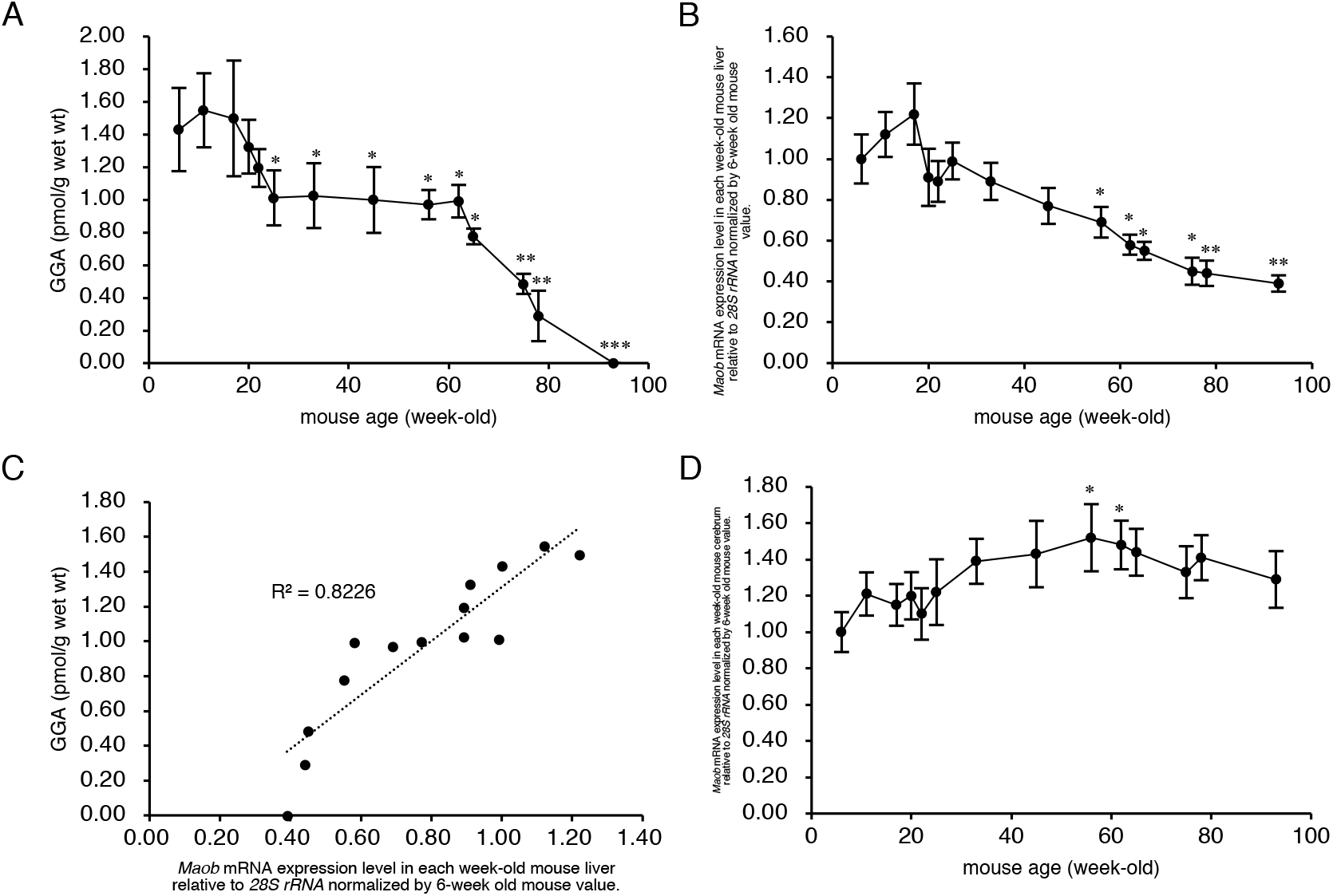
Association of age-dependent changes in hepatic GGA content with *Maob* mRNA level in the liver of C3H/HeN mice. A: Endogenous GGA contents in the liver of each week-old male C3H/HeN mouse. B: The relative expression levels of *Maob* mRNA in each week-old mouse liver to 6-week old mouse liver. C: Correlation between the amount of endogenous GGA and expression level of *Maob* mRNA in C3H/HeN mouse liver. Each point represents the average value of each age group (n = 5). D: The relative expression levels of *Maob* mRNA in each week-old mouse cerebrum to 6-week old mouse cerebrum. All points represent the mean ± SEM (*n* = 5). *; *P* < 0.05, **; *P* < 0.01 and ***; *P* < 0.001 versus 6-week old mice (ANOVA with post hoc Scheffe). GGA, geranylgeranoic acid; *Maob*, monoamine oxidase B

### *Maob* expression in C3H/HeN mouse liver decreases with aging

Next, we investigated whether the expression level of *Maob* mRNA, which was reported in our previous study to be involved in the biosynthesis of endogenous GGA, is responsible for the age-dependent decrease in hepatic GGA level. *Maob* mRNA expression levels in mouse liver started to decrease gradually and constantly from 18 weeks of age, and significantly decreased after 56-week old compared to 6-week old mice [**Figure 3B**]. Hence, there was a significant and positive correlation between endogenous GGA level and *Maob* mRNA level in male C3H/HeN mouse liver [**Figure 3C**]. The expression level of *Maob* mRNA in C3H/HeN mouse cerebrum increased inversely with aging [**Figure 3D**]. *Maob* mRNA expression levels between the liver and cerebrum in C3H/HeN mice indicated a weak negative correlation [see **Supplementary Figure 2A**].

On the other hand, mRNA expression level of the *Maoa* gene, another member of the flavin monoamine oxidase family, gently increased along with aging in C3H/HeN mouse liver [**Figure 4A**], so that there was an apparent negative correlation between *Maoa* mRNA expression level and the amount of GGA in the liver [**Figure 4B**]. The age-related variation in *Maoa* mRNA levels in the cerebrum was a mirror image of the age-related variation in *Maob* mRNA levels in the cerebrum [**Figure 4C**]. *Maoa* mRNA expression levels between the liver and cerebrum in C3H/HeN mice indicated a weak positive correlation [see **Supplementary Figure 2B**].

**Figure 4.**
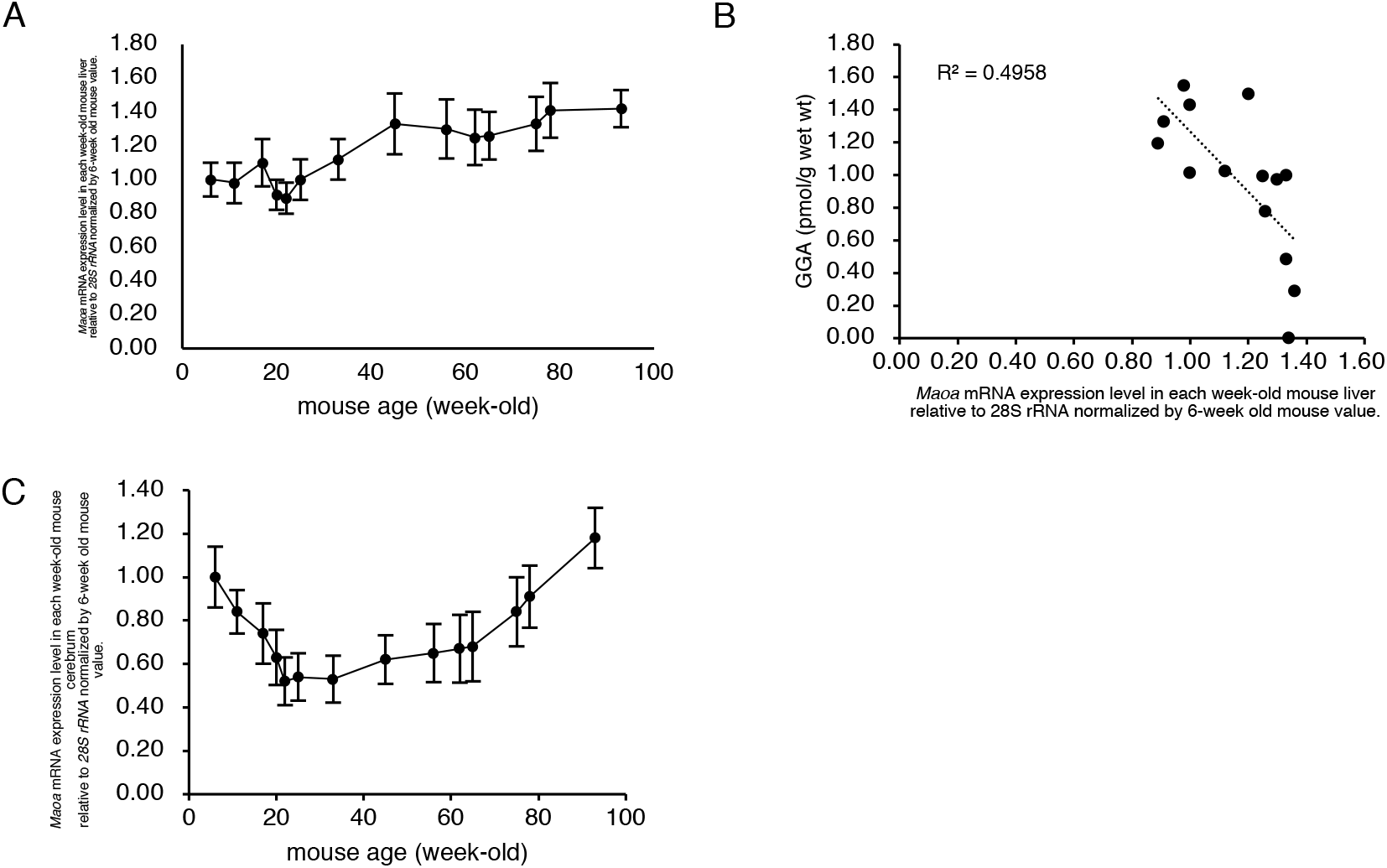
Apparent inverse correlation of hepatic *Maoa* mRNA levels and hepatic GGA content in C3H/HeN mice. A: The relative expression levels of *Maoa* mRNA in each week-old mouse liver to 6-week old mouse liver. B: Correlation between the amount of endogenous GGA and hepatic expression level of *Maoa* mRNA. Each point represents the average value of each age group (n = 5). C: The relative expression levels of *Maoa* mRNA to 6-week old mouse cerebrum in each week-old mouse cerebrum. All points represent the mean ± SEM (*n* = 5). GGA, geranylgeranoic acid; *Maoa*, monoamine oxidase A

### *Maob* mRNA expression is further reduced in cancerous liver tissue

Since age-related decreases in GGA and *Maob* mRNA expression levels were observed in the normal liver of C3H/HeN mice, next, we measured GGA, *Maob* mRNA, and *Maoa* mRNA levels in the tissue of spontaneously developed mouse hepatoma in comparison with those in its surrounding normal liver tissue [see **Supplementary Figure 3**]. In 94-week old mice, *Maob* mRNA expression level was further and significantly reduced in hepatoma tissue compared to that of its surrounding normal liver tissue, which was 40% of hepatic *Maob* mRNA level of 6-week old mice [**Figure 5A**]. Whereas, there was no significant difference in *Maoa* mRNA expression level between hepatoma tissue and the normal liver tissue of 94-week old mice [**Figure 5B**].

**Figure 5.**
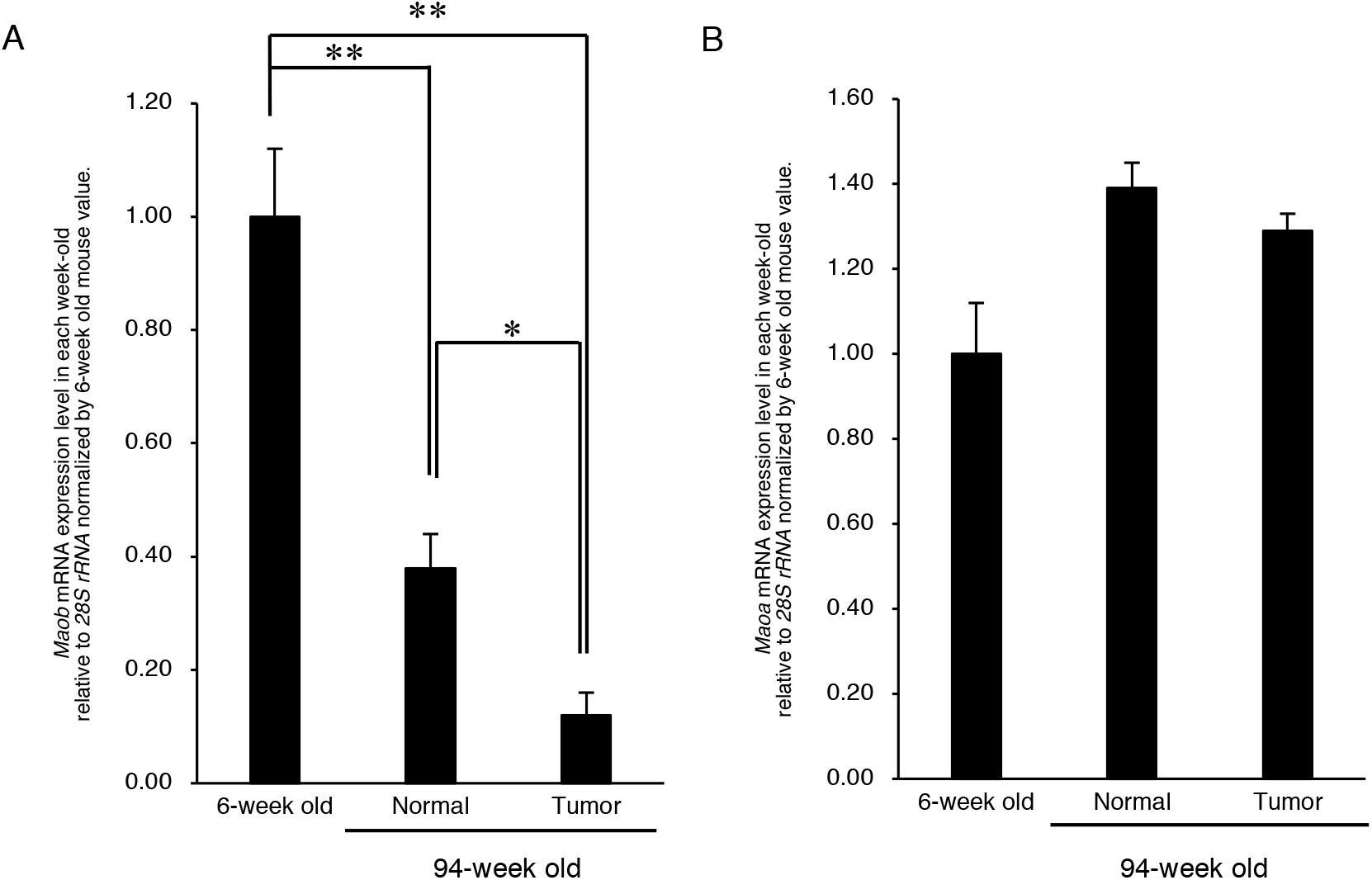
*Maob* mRNA and *Maoa* mRNA levels in liver tumor tissue and its surrounding normal tissues. The relative expression levels of *Maob* mRNA (A) and *Maoa* mRNA (B) in the liver tumor and its surrounding normal hepatic tissues of 94-week old C3H/HeN mice liver. The relative expression levels of each mRNA were expressed as compared with those of each mRNA in 6-week old mouse liver. The data represent the mean ± SEM (*n* = 5). *; *P* < 0.05 and **; *P* < 0.01 versus 6-week old mice (ANOVA with post hoc Scheffe). *Maoa*, monoamine oxidase A; *Maob*, monoamine oxidase B;

### A single dosing of GGA to 11-month-old C3H/HeN mice can efficiently suppress the development of spontaneous hepatomas

Assuming that the age-related decline in liver GGA content observed above is permissive for the spontaneous development of hepatoma, we hypothesized that supplementation with GGA at the time of the decline in liver GGA would prevent the spontaneous development of hepatoma. In this context, we examined whether administration of GGA to mice at an appropriate time would prevent spontaneous hepatocarcinogenesis in male C3H/HeN mice.

Our result clearly shows a significant relationship between the timing of a single administration and the hepatoma-preventive effects of GGA. In other words, a single oral supplementation with GGA at 11 months of age most strongly inhibited spontaneous hepatocarcinogenesis at 23-month old [**Figure 6**]. In more detail, the average number and weight of 23-month-old mouse liver tumors were most significantly suppressed to approximately 30% in the group administered GGA at the age of 11 months compared to the control group. When GGA was administered to mice at 7 months of age, the average number and weight of spontaneous hepatomas per mouse were reduced by roughly 40% compared to those of hepatomas in the control mice at 23-month old, but the effects of GGA were not statistically significant. GGA administration at 17-month old also reduced the average weight of liver tumors by approximately 40%, but the average number of liver tumors per mouse was not reduced.

**Figure 6.**
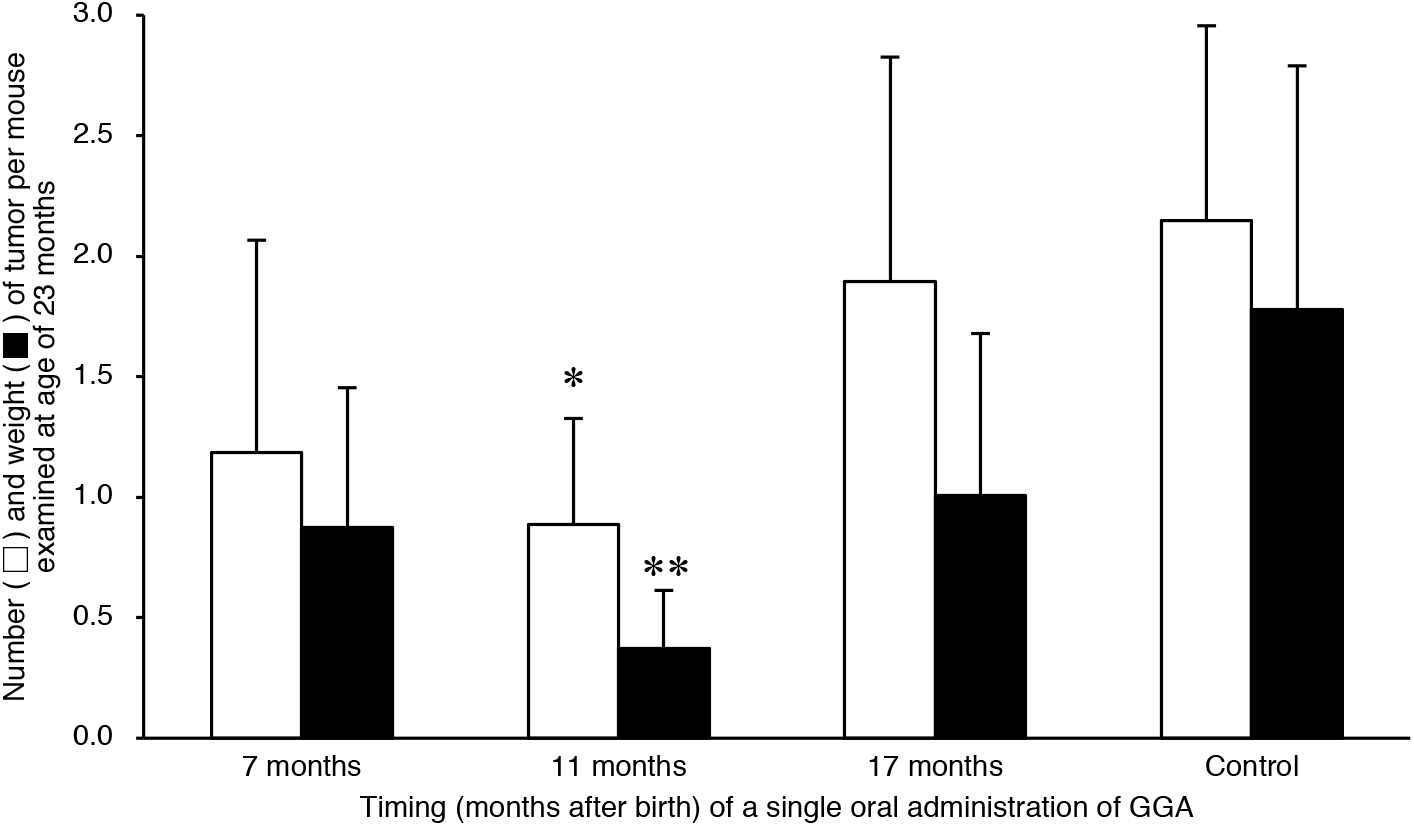
Relationship between timing of a single dosing of GGA and inhibitory effect on tumor development in C3H/HeN mice. The number and weight (g) of liver tumors at 23-month old in the group of mice to which GGA was orally administered only once at age of 7, 11, or 17 months are shown. Control is the group without orally administration of GGA. The data represent the mean ± SD (n = 10). \**P* < 0.05 and \*\**P* < 0.02 (versus control mice, ANOVA with post hoc Scheffe). GGA, geranylgeranoic acid;

### Dietary intake of GGA or GGOH increases hepatic GGA level of C3H/HeN mice

Finally, we examined a couple of ways to increase hepatic GGA content of C3H/HeN mice. Firstly, we tried to induce an increase in hepatic GGA level of C3H/HeN mice by dietary intake of exogenous GGA. Dietary intake of GGA for a week dose-dependently increased hepatic GGA level in 24-week old mice [**Figure 7A**]. On the other hand, dietary GGA did not significantly change hepatic free ARA concentration [see **Supplementary Figure 4**]. Secondly, GGOH, a metabolic precursor of GGA and a substrate of MAOB was examined. Dietary intake of GGOH for a week dose-dependently increased hepatic GGA level up to a dose of 4 mg/day, but no further increase was observed with dietary GGOH at a dose of 8 mg/day [**Figure 7B**]. Dietary GGOH also did not significantly change hepatic ARA level (data not shown).

**Figure 7.**
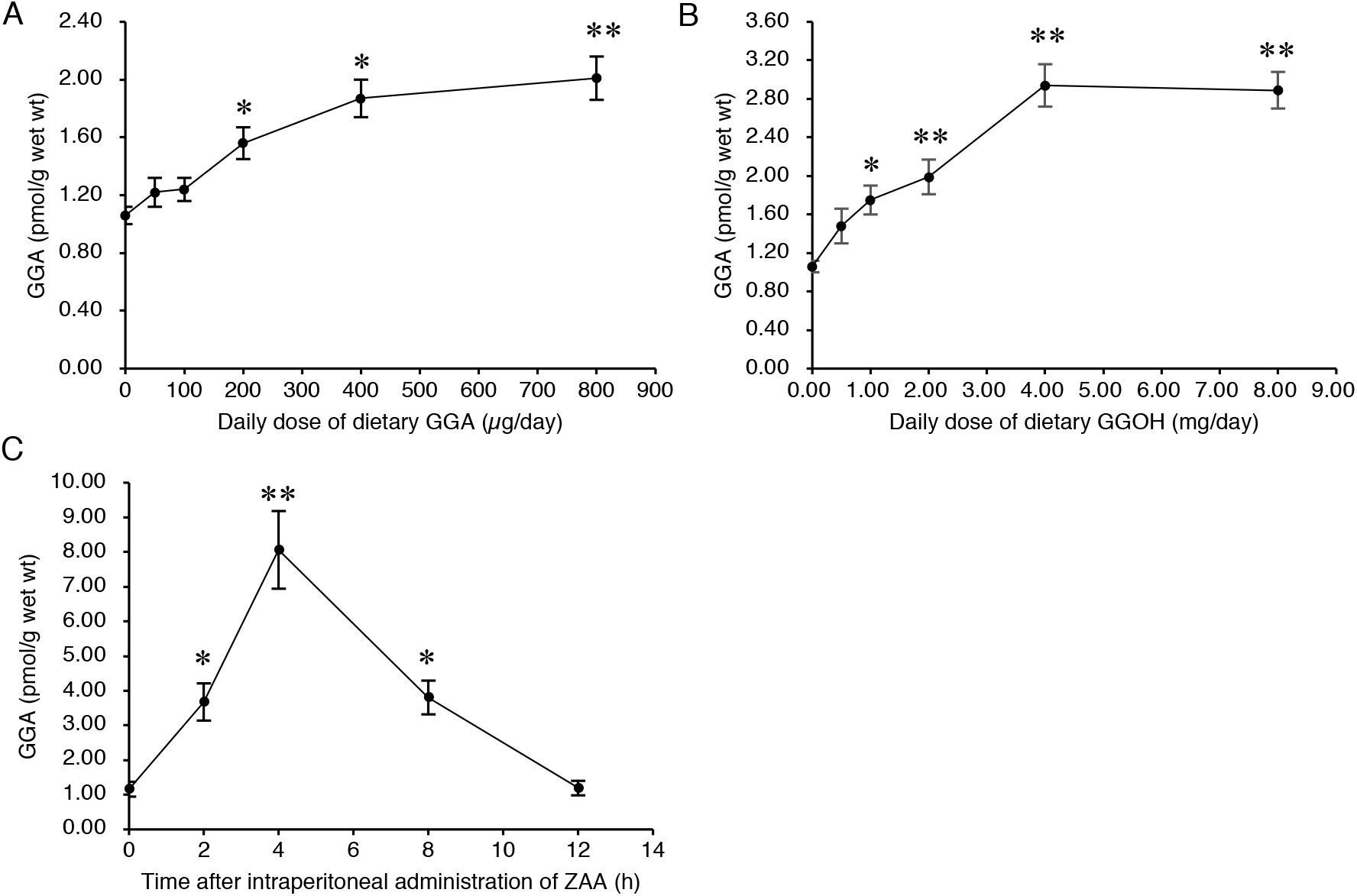
Upregulation of hepatic GGA level by either oral administration of GGA or GGOH or intraperitoneal administration of ZAA. The content of GGA in the 24-week old C3H/HeN mouse liver after 1-week oral administration of GGA (0–800 μg/day/mouse) (A) or GGOH (0–8 mg/day/mouse) (B). Time course of hepatic GGA contents in the 24-week-old C3H/HeN mice after the intraperitoneal administration of ZAA (C). The results are expressed as the mean ± SD (*n* = 3). \**P* < 0.05 and \*\**P* < 0.01 versus mice fed a control diet (A and B) or versus untreated mice (C). (ANOVA with post hoc Scheffe). GGA, geranylgeranoic acid; GGOH, geranylgeraniol; ZAA, Zaragozic acid A

### Intraperitoneal administration of ZAA upregulates hepatic GGA level of C3H/HeN mice

Thirdly, intraperitoneal administration of ZAA was performed to upregulate hepatic GGA level of the C3H/HeN mice without any dietary manipulation. Similar to ZAA treatment of human hepatoma-derived cell line^[17]^, GGA levels in the liver increased transiently from a baseline of 1.16 to a peak of 8.06 pmol/g at 4 h after the administration of ZAA. The GGA level in the liver increased by ZAA returned to the initial level after 12 h of administration [**Figure 7C**]. Similar to dietary GGA and GGOH, intraperitoneal administration of ZAA also did not significantly change hepatic ARA level (data not shown).

## DISCUSSION

In the present study, we examined the hypothesis whether reduced hepatic GGA level may allow spontaneous development of hepatomas in male C3H/HeN mice. To this end, we first demonstrated an age-dependent decline of hepatic GGA level. A similar age-dependent decrease in hepatic *Maob* mRNA levels was also observed, resulting in a strong positive correlation between liver endogenous GGA and hepatic *Maob* mRNA levels. In addition, we found that *Maob* mRNA expression level was significantly reduced in spontaneous liver tumor tissue compared to its surrounding normal liver tissue in 94-week-old male C3H/HeN mice. Next, we were able to successfully suppress spontaneous hepatoma development at 23 months of age by a single oral dosing of GGA (50 μg) to 11-month-old male mice. These results indicate that the age-dependent decline of hepatic GGA level is potentially permissive with spontaneous hepatocarcinogenesis in male C3H/HeN mice and that hepatic carcinogenesis can be prevented by some upregulation of hepatic GGA level.

In the course of our study on hepatoma prevention, initially, acyclic retinoids of GGA and 4,5-didehydroGGA were synthesized chemically as a preventive agent against a second primary hepatoma and we found that GGA and 4,5-didehydroGGA induced cell death in several human hepatoma-derived cell lines, but not in mouse primary hepatocytes^[8,9,14]^. Afterwards, we reported that GGA is a natural compound found in medicinal plants such as turmeric^[14]^, but, recently, we reported using human hepatoma-derived HuH-7 cells that GGA is also an endogenous lipid that is enzymatically biosynthesized through the mammalian MVA pathway, and demonstrated that ZAA treatment can induce cell death in HuH-7 cells by increasing intracellular GGA^[17,18]^. Hence, we proposed that when intracellular GGA is above a certain concentration, hepatocytes in the precancerous stage become GGA-sensitive at a certain point in the carcinogenesis process, leading to cell death, which in turn eliminates hepatocarcinogenic cells. Following this hypothesis, we can speculate that in C3H/HeN mice that spontaneously develop hepatomas, hepatic GGA levels would decrease with age, which would allow spontaneous hepatoma to develop at 2 years of age. Thus, in the present study, we investigated whether hepatic GGA level in C3H/HeN mice was decreased before hepatocarcinogenesis. First, we measured endogenous GGA content in the liver of the young mice. As a result, we found that hepatic GGA level in 6-week-old male C3H/HeN mice was 1.4–1.6 pmol/g, which is considerably lower than the liver GGA level of 80 pmol/g in 5-week-old male Wistar rats^[17]^ and even the endogenous GGA levels of human hepatoma-derived HuH-7 (10–25 pmol/g)^[25]^ and Hep3B (10–15 pmol/g)^[18]^ cells.

One of the most important findings of the present study is the age-related variation in liver GGA concentrations in male C3H/HeN mice, showing that hepatic GGA levels decrease in two age-dependent stages. The first stage corresponds to 20 to 25 weeks of age and hepatic GGA level decreases rapidly from 1.5 to 1.0 pmol/g, but then remains constant at 1.0 pmol/g until 60 weeks. Then, in the second stage, there is a linear decrease from 60 to 90 weeks, and at 93 weeks, hepatic GGA level reaches below the detection limit. Since age-related changes in total liver GGA levels (pmol/liver/mouse) also showed a two-step decrease similar to hepatic GGA concentrations (pmol/g liver) described above [see **Supplementary Figure 5**], indicating that the two-step decrease in hepatic GGA levels due to aging is not due to changes in liver organ weight. Considering that there was no change in liver weight of mouse in 62 to 93-wk old (1.12–1.19 g), the decrease in hepatic GGA level with these ages is not due to a dilution of hepatic GGA level by hepatic hypertrophy, but may be due to putative metabolic declines of hepatic GGA. Not surprisingly, GGA was undetectable in several spontaneous hepatomas or in any of the surrounding normal areas at week 94 (unpublished results). Unfortunately, in the present experiments in which the liver was analyzed over time [Figure 3], we could not visually detect enough to dissect hepatoma areas in mice before 90 weeks of age, so we cannot describe the difference in GGA levels between cancerous and non-cancerous areas before hepatic GGA was completely depleted. But, we are sure that GGA is also depleted in normal liver tissue around liver tumor tissue. Taking into account the high recurrence (strictly speaking, second primary tumor) rate in the remaining normal liver tissue after radical therapies to eliminate human liver tumor^[26]^, a decrease in hepatic GGA content may predispose hepatocytes to hepatoma development, which may be regarded as one of the new causes of “field cancerization” in the liver^[27]^.

GGA is a C_20:4_ polyunsaturated fatty acid. In C3H/HeN mice, hepatic contents of free ARA, an endogenous lipid that is a structural isomer of GGA, did not decrease with aging, but rather increased in tumor tissue (unpublished results). The increase in ARA in tumor tissue observed here is consistent with previous studies showing that ARA acts as an inflammatory mediator via eicosanoids^[28,29]^. In other words, it can be considered hepatic fatty acid synthesis does not decrease in general with aging of C3H/HeN mice, and endogenous GGA levels specifically decreased in C3H/HeN mouse liver, which is probably due to a decrease in the GGA biosynthesis. Of note, this is consistent with our previous finding that a mirror-image relationship between endogenous ARA and endogenous GGA levels has been detected in different growth states of human hepatoma-derived cell lines^[25]^.

We have recently shown by knockdown experiments that intracellular GGA levels positively correlate well with *MAOB* mRNA expression levels in human-hepatoma derived HuH-7 and Hep3B cells^[18]^. Here we have demonstrated an age-dependent decrease in hepatic *Maob* mRNA expression levels, which may be one of the reasons for the age-dependent decrease in hepatic GGA contents of male C3H/HeN mice. Indeed, *Maob* mRNA expression level significantly and positively correlated with the endogenous GGA content, suggesting that hepatic MAOB is the rate-limiting enzyme for GGA synthesis. In contrast to *Maob* mRNA, *Maoa* mRNA expression levels tended to increase with age, resulting in strong and negative correlation of *Maoa* mRNA levels with endogenous GGA content in the liver. On the other hand, similar to hepatic *Maoa* mRNA, the expression levels of the cerebrum *Maob* and *Maoa* mRNAs in C3H/HeN mice increased with age. These age-dependent changes of *Maob/Maoa* expression levels in male C3H/HeN mouse liver and brain are consistent with age-dependent changes of MAOB and MAOA protein levels in male Bl/C57 mouse liver and brain reported by Saura *et al*^[20]^, respectively.

Although no GGA was detected in the hepatoma area or in its surrounding normal tissues of 94-week-old mice, it is still worth considering that the expression levels of *Maob* mRNA was significantly lower in the hepatoma area than in the surrounding tissues [Figure 5A]. Up to the present, there have been several reports of reduced MAO activity in the hepatoma area, which is considered MAOB activity. White and Tewari reported that mitochondrial benzylamine oxidase (equivalent to MAOB) activity was null in Novikoff hepatoma transplanted in rats, whereas other mitochondrial oxidoreductases including tryptamine oxidase (equivalent to MAOA) remained considerable activity in the transplanted hepatoma^[30]^. Of note, using Morris hepatoma cell lines of varying degrees of differentiation, Pedersen *et al*. showed that benzylamine oxidase activity decreased with decreasing degrees of differentiation^[31]^. And they speculated that the synthesis of “MAOB” might be specifically repressed in hepatoma and proposed that a deficiency of “MAOB” activity in mitochondria is a reflection of an early stage of the carcinogenic process^[31]^. Taken together with these classical findings and our present results, we speculate that the age-dependent decrease of *Maob* mRNA in C3H/HeN mouse liver results in a decrease in MAOB activity with age, leading the liver to the early stages of the carcinogenic process. These classical studies proposed that enzymatic degradation of some carcinogenic amines is attenuated by reduced MAOB activity, but it is unlikely that abnormal accumulation of carcinogenic amines is the cause of spontaneous hepatocarcinogenesis in C3H/HeN mice. Here, we can reasonably speculate that the decrease in hepatic MAOB activity induces a decrease in endogenous GGA levels in the liver, allowing cells in the carcinogenic stage to escape GGA-induced cell death and leading to the development of spontaneous hepatomas. In other words, the emergent escape and survival from death signals, and later outgrowth of malignant cells, should be at least partially attributed to a deficiency in cellular GGA content due to downregulation of MAOB in the early stages of hepatocarcinogenesis. Since MAOB-expressing premalignant cells should not have decreased cellular GGA content, even if such cells are present, they will undergo cell death via GGA-induced pyroptosis, leading to spontaneous prevention of hepatocarcinogenesis^[32]^. Indeed, although it is not hepatoma, the possible preventive roles of MAOB activity in carcinogenesis are further supported by experiments using enzyme inhibitors^[33]^: prolonged administration of a MAOB inhibitor, but not a MAOA inhibitor, significantly increased colon carcinogenesis induced by azoxymethane.

In this context, next, we investigated whether oral supplementation of GGA suppresses hepatocarcinogenesis when GGA in the liver of C3H/HeN mouse is decreased. In doing so, we referred to Watanabe & Omori’s pioneering work^[34]^, which described that a single dosing of 4,5-didehydroGGA (peretinoin) at the age of 12 months was most effective to suppress spontaneous hepatomas in male C3H/HeN mice at the age of 23 months. Here, we also found that a single administration of GGA was most effective at approximately 11 months after birth to prevent hepatocarcinogenesis, but less effective at 7 months and ineffective at 17 months of age. A quarter century ago, we reported that a single administration of 4,5-didehydroGGA to 12-month-old C3H/HeN mice produced the most prominent anti-oncogenic effect at the age of 23 months, but the reason for this was unknown at the time^[35]^. But, now, we are able to speculate that a narrow window at approximately 11–12 months after birth is a timing when hepatic GGA content age-dependently decreases to a critical level that loses its ability to prevent carcinogenesis. A single dose of GGA to 11-month-old mice would remove from the body all carcinogenic cells that had escaped the cell death due to a decreased endogenous GGA, thus eliminating the possibility of carcinogenesis at 23 months of age. It is understandable that the preventive effect of early administration of GGA is weak because the GGA level in the liver does not decrease until 6 months of age. Moreover, it is also possible that precancerous hepatocytes that have acquired GGA sensitivity might not have developed during this period. On the other hand, because hepatic GGA declines rapidly after 17 months of age and is depleted by 23 months of age, administration of GGA during this period may be expected to be effective. However, the number of hepatomas at 23 months of age in mice treated with a single dose of GGA at 17 months of age was not different from that of the untreated controls, but the weight of hepatoma per mouse was significantly reduced. At the present moment, we are speculating that at 17 months, carcinogenesis should be in its complete stage, and cancerous tissues may have already become resistant to GGA-induced pyroptotic cell death.

Finally, we examined dietary and pharmacological methods to increase hepatic GGA levels in C3H/HeN mice. Previously, we reported that GGA levels in human serum increased by oral ingestion of turmeric tablets containing natural GGA^[36]^. And, since we recently found that adding GGA to the diet during breeding of C3H/HeN mice effectively increased the fertility^[37]^, we decided to first examine whether GGA added to the diet would increase hepatic GGA content. As expected, the addition of GGA to the diet increased hepatic GGA levels in a dose-dependent manner, and at a dose of 400 μg/day, hepatic GGA level was nearly saturated, almost double the normal value [Figure 7A]. However, the increase in hepatic GGA was only 20% at a daily dose of 50 μg, which is the effective dose in the single-dose carcinogenesis suppression experiments, and therefore the kinetics of GGA in the liver after a single dose of GGA should be examined in the future.

We have previously shown that GGOH produced from geranylgeranyl diphosphate by intestinal alkaline phosphatase is converted to GGA by liver enzymes, suggesting that GGOH can be used as an exogenous source of hepatic GGA^[38]^. The dose was about 10 times greater than GGA, but in a dose-dependent manner exogenous GGOH increased hepatic GGA levels, reaching 2.8 times the normal value at a dose of 4 mg/day and saturating [Figure 7B]. Considering the availability and price from the market, GGOH would be a more efficient candidate for raising liver GGA levels than GGA itself. However, compared to GGA, GGOH should be taken in larger doses, and the possible side effects of GGOH may need to be studied in the future. Of great interest to us, oral administration of GGOH to rats suppresses the early stages of hepatocarcinogenesis^[39]^. The present results allow us to hypothesize that GGOH is metabolized to GGA in the livers of GGOH-treated rats, and the resultant GGA may inhibit the development of liver tumors.

In addition, besides the nutritional methods mentioned above, pharmacological approaches are also possible to increase hepatic GGA levels. As an example, we performed intraperitoneal administration of ZAA to mice, because the addition of ZAA to the human hepatoma-derived HuH-7 cell culture system increased endogenous GGA levels and induced cell death^[17]^. At 4 h after intraperitoneal administration to C3H/HeN mice, ZAA increased hepatic GGA levels up to eight times the pre-injection level, but returned to the original value after 12 h, indicating that ZAA is metabolized as a drug and is rapidly deactivated. Therefore, it may be possible to take ZAA only during times of high risk for hepatocarcinogenesis, such as after radical therapy of hepatoma or cirrhosis, to wipe out the carcinogenic cells from the body, resulting in prevention of second primary hepatomas.

In any case, to establish the chemoprevention of hepatoma by dietary GGA/GGOH or a drug of ZAA in humans, we absolutely need to elucidate more detailed molecular biological mechanism of hepatocarcinogenesis surveillance mechanism by endogenous GGA in the liver.

In conclusion, we showed that both GGA and *Maob* mRNA expression levels in the male C3H/HeN mouse liver were significantly reduced at the pre- or early stage of liver carcinogenesis, and that oral supplementation of GGA prevents hepatocarcinogenesis. In the future, our research offers the potential for chemical prevention of hepatoma with dietary or pharmacological manipulation of hepatic GGA contents.

## Abbreviation

ARA: arachidonic acid
ATRA: all-*trans* retinoic acid
GGOH: geranylgeraniol
GGA: geranylgeranoic acid
GSDMD: gasdermin D
MVA: mevalonic acid
MAOA: monoamine oxidase A
MAOB: monoamine oxidase B
TLR4: Toll-like receptor 4
ZAA: zaragozic acid A

## DECLARATIONS

## Acknowledgments

The authors are extremely grateful to Yukiko Araki, Asaka Muraoka, Sayaka Uematsu, Reina Arakaki, Rina Noaki, Maho Hattori, Yoshika Yasui, and Miyuki Yano of their laboratory members who dedicated a lot of time for mice caring.

## Authors’ contributions

Made contributions to original conceptualization: Omori M, Shidoji Y;

Made contributions to experimental design for original idea: Omori M, Shidoji Y, Tabata Y;

Made contributions to performed data acquisition, analysis and interpretation: Tabata Y, Omori M, Shidoji Y;

Made contributions to design, writing and revision of the paper: Tabata Y, Shidoji Y.

## Ethical approval and consent to participate

Not applicable.

