## Supplementary Table or Supplementary Figure for "Age-dependent decrease of hepatic geranylgeranoic acid and its oral supplementation prevents spontaneous hepatoma in C3H/HeN mice"

**Supplementary Table 1. The nucleotide sequences of each primer used for RT-qPCR.**

| Genes | Primer | Sequence (5’ – 3’) |
|---|---|---|
| *Maoa* | F | GGTCCTCCTTGGGGATAAAG |
| | R | TTCTCTCAGGTGGAAGCTCTG |
| *Maob* | F | GCACTGAAACAGCCTCACAC |
| | R | TCGTGCAGGGACATCCAAAG |
| *28S rRNA* | F | GCTCAGTACGAGAGGAACCG |
| | R | AGAGGCGTTCAGTCATAATC |

F: forward primer, R: reverse primer

*Maoa*, monoamine oxidase a; *Maob*, monoamine oxidase b.

**Supplementary Table 2. The condition of thermal cycler for RT-qPCR of *Maoa*, *Maob and 28s rRNA*.**

| | Temperature, Duration | Slope |
|---|---|---|
| Denature | 95ºC, 600 s | 20ºC / s |
| PCR (40 cycles) | 95ºC, 15 s | 20ºC / s |
| | 65ºC, 60 s | 20ºC / s |
| Melting | 95ºC, 0 s | 20ºC / s |
| | 57ºC, 15 s | 20ºC / s |
| | 98ºC, 0 s | - |
| Cooling | 40ºC, 30 s | 20ºC / s |

*Maoa*, monoamine oxidase a; *Maob*, monoamine oxidase b.

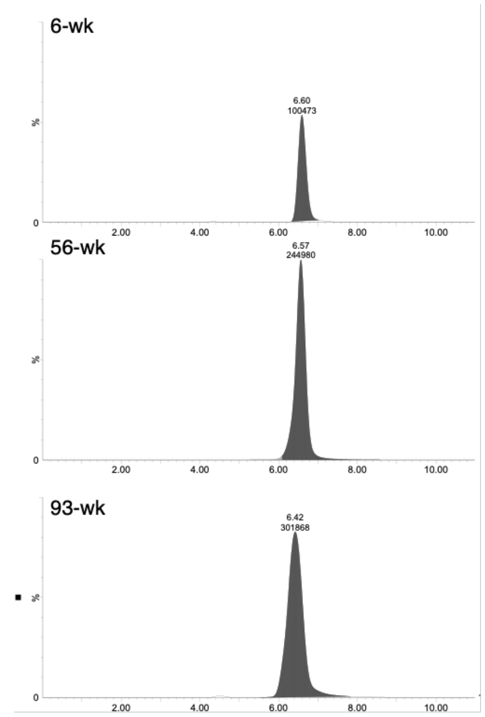


**Supplementary Figure 1.** Representative LC/MS/MS chromatograms of the 6-, 56- and 93-week old mouse hepatic lipid extracts tracing arachidonic acid (m/z: 303-259). The chromatograms show a peak height of $7.87 \times 10^5$ as 100%.

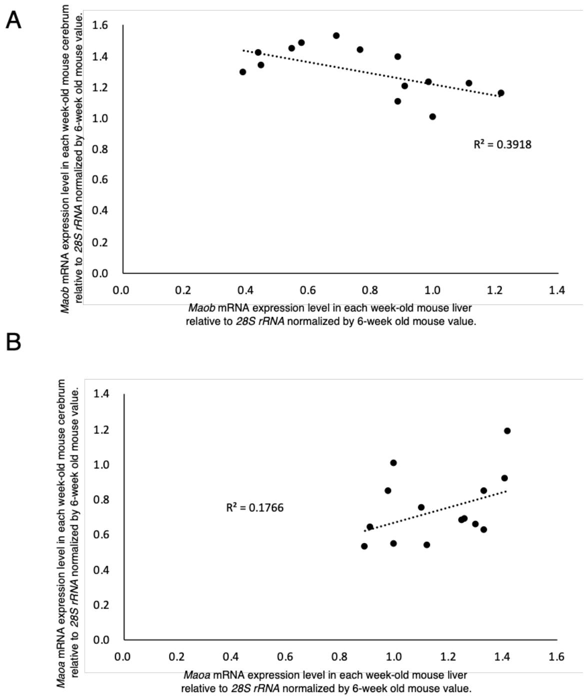


**Supplementary Figure 2.** A: Association of hepatic *Maob* mRNA with cerebrum *Maob* mRNA levels. B: Association of hepatic *Maoa* mRNA with cerebrum *Maoa* mRNA levels. Each point represents the average value of each age group ($n = 5$). *Maoa*, monoamine oxidase a; *Maob*, monoamine oxidase b.

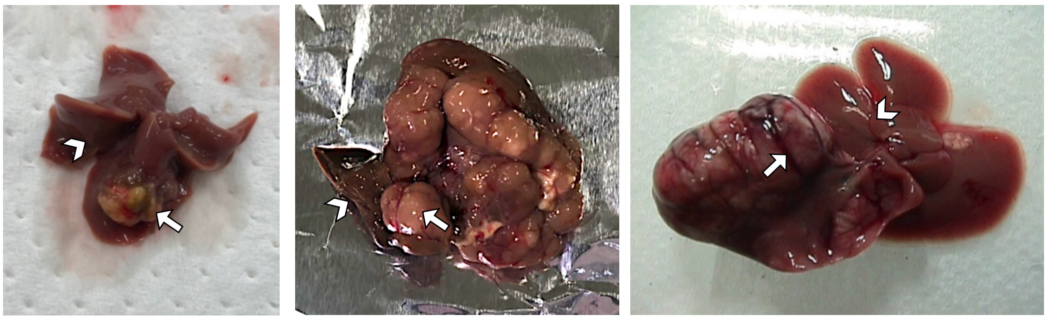

**Supplementary Figure 3.** Representative normal tissue (arrowhead) and tumor tissue (arrow) of male C3H/HeN mice used for measurement of endogenous GGA and *Maoa/Maob* mRNA levels.

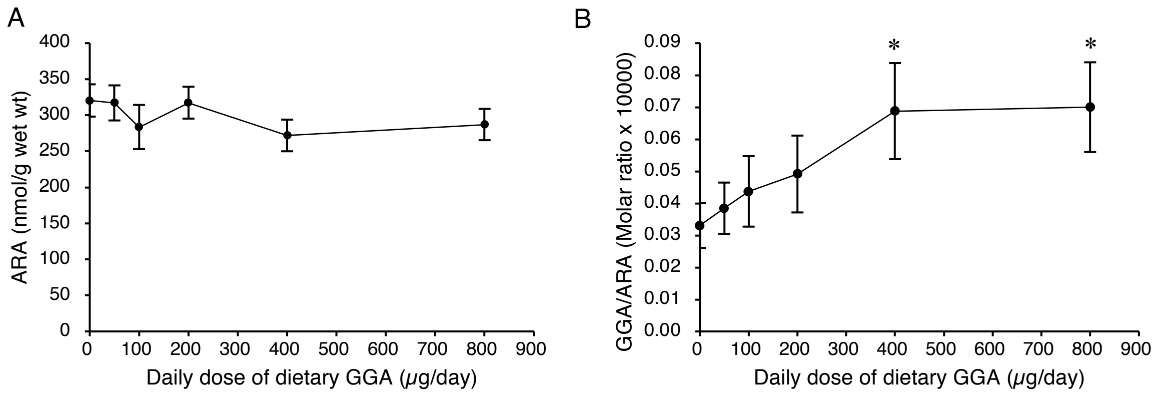


**Supplementary Figure 4.** Hepatic free ARA levels (A) and GGA/ARA molar ratio (B) in mice liver are plotted along with daily dose of orally administered GGA. The results are expressed as the mean ± SD ($n$ = 3). *$P < 0.05$ versus mice fed a control diet (ANOVA with post hoc Scheffe). ARA, arachidonic acid; GGA, geranylgeranoic acid;

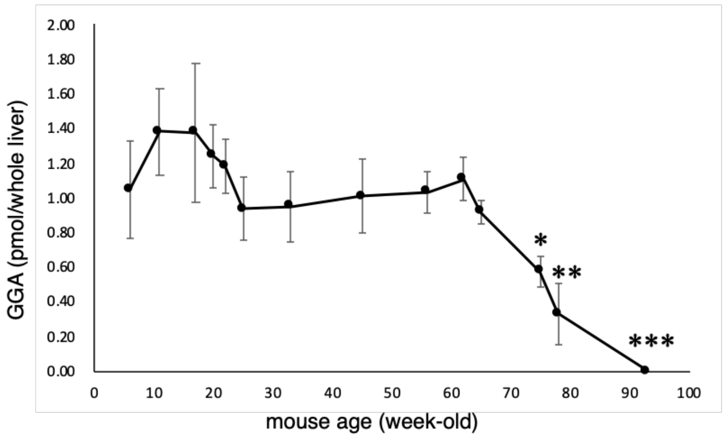


**Supplementary Figure 5.** Age-dependent changes of hepatic GGA content per C3H/HeN mouse. Endogenous GGA contents in the whole liver of each week-old male C3H/HeN mouse are plotted. All points represent the mean ± SEM ($n$ = 5). *$P < 0.05$, **$P < 0.01$ and ***$P < 0.001$ versus 6-week old mice (ANOVA with post hoc Scheffe). GGA, geranylgeranoic acid.
